# Gene-First Identity Construction for Robust Cell Identification in Single-Cell Transcriptomics

**DOI:** 10.64898/2026.02.25.707869

**Authors:** Luqi Yang, Zhenwei Huang, Jinpu Cai, Hongyi Xin

## Abstract

The precise delineation of cell types is fundamental to single-cell transcriptomics, yet current clustering pipelines often violate an axiomatic principle: hierarchical consistency. Existing methods measure cell-to-cell distances within a fixed global feature space, disregarding the fact that biological distinctions are inherently context-dependent lineage separation requires different gene programs than subtype resolution. Mathematically, this implies that the similarity metric itself should not be a static functional, but a pair-dependent energy functional evaluated within a specific Hilbert subspace determined by the biological comparison at hand. The challenge lies in the fact that allowing pair-dependent metrics typically destroys the global geometric consistency required for downstream analysis, unless the family of Hilbert subspaces is given strong biological structure. To resolve this geometric dilemma, we introduce GeCCo (Gene Co-expression Constructed identity), which constructs identities by projecting cells onto a rigorously derived hierarchy of gene programs. To construct this hierarchy, GeCCo first quantifies Boolean regulatory logic via the *ϕ* coefficient, and subsequently employs a greedy topological inference to organize genes based on their synergistic and antagonistic relationships. Benchmarking on human immune atlases demonstrates that GeCCo achieves superior hierarchical consistency, ensuring that globally inferred cell identities rigorously match locally refined subtypes. Furthermore, in pancreatic endocrine progenitors, GeCCo resolves a hidden mitotic bridge state, suggesting a concentrated division phase prior to differentiation. Ultimately, GeCCo shifts the paradigm from ad hoc clustering to programmatic cell typing, offering a mathematically grounded framework for scalable atlases of cellular discovery.

## 1 Introduction

In the era of large-scale single-cell atlases, accurate and stable cell-type definition has become a central challenge in computational biology[1, 2]. Projects such as the Human Cell Atlas[3], Tabula Sapiens[4], and Mouse Cell Atlas[5] encompass millions of cells across diverse tissues, developmental stages, and species. This scale has made the field increasingly reliant on cell identity systems that are both consistent and reproducible. Critically, cell identity is not a flat labeling scheme but a naturally hierarchical structure, spanning broad lineages, fine-grained subtypes, and transient transitional states[6]. Consequently, computational methods designed to recover this hierarchy robustly and biologically are essential for reliable cross-dataset integration and mechanistic interpretation.

However, current gold-standard analytical pipelines, such as Seurat[7] and Scanpy[8], face a fundamental limitation when resolving this multi-scale structure: *hierarchical inconsistency*. These methods typically rely on the identification of Highly Variable Genes (HVGs) across the input population to drive clustering. While effective for capturing global heterogeneity, this strategy often yields inconsistent subclusters when applied globally versus locally to the same set of cells[9]. This phenomenon is explicitly quantified in Figure 1a–c, using a dataset comprising three major lineages (A, B, C) and their respective subtypes. We first established a biologically consistent reference by independently sub-clustering each lineage (Fig. 1a) and aggregating the resulting labels (Fig. 1b). In contrast, when clustering was performed directly on the full dataset (Fig. 1c), the resulting partition (*k* = 6) deviated significantly from this reference structure, yielding a low Adjusted Rand Index (ARI = 0.09). This inconsistency arises not from the clustering algorithms themselves, but from the context-dependency of feature selection. As detailed in Supplementary Figure S1, feature selection prioritizes different sets of genes depending on the analysis scope: markers driving local subtype distinctions are often overshadowed by macro-lineage markers in a global context. Consequently, global PCA extracts principal components reflecting broad lineage structures, while missing the distinct axes of variation necessary to resolve subtle subtypes. Ultimately, if cell identities shift depending on the analytical starting point, the resulting partitions are computationally unstable and biologically unreliable.

**Fig. 1.**
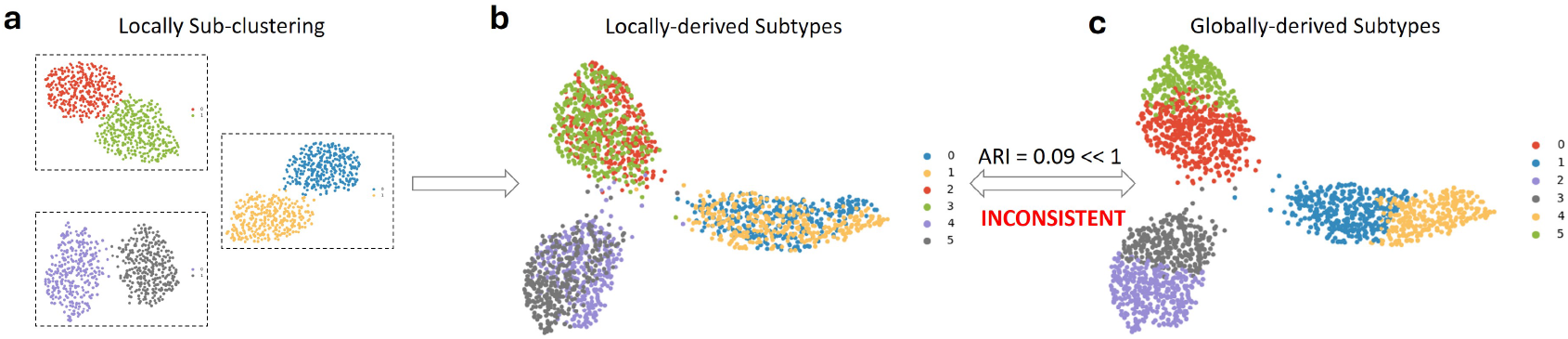
Hierarchical inconsistency arising from global context bias. . **(a)** The local clustering: Major lineages (A, B, C) are extracted and clustered independently to obtain subtype labels. **(b)** Global UMAP colored by the *aggregated* labels derived from the local process in (a). **(c)** Global UMAP colored by labels derived from *direct global clustering* of the entire dataset. Note the severe mismatch (ARI = 0.09) compared to the aggregated local labels.

**Fig. 2.**
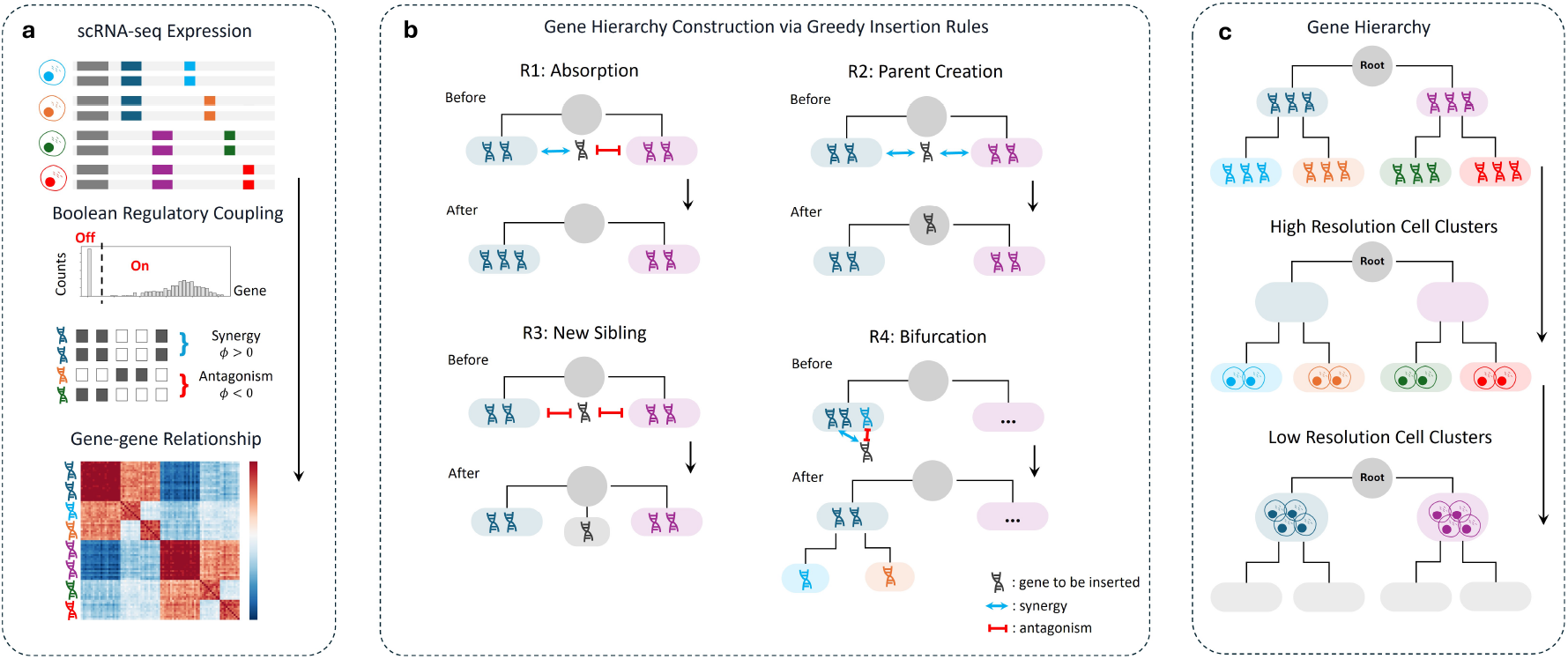
Schematic overview of GeCCo framework. **(a)** Raw scRNA-seq expression matrices are projected onto a boolean regulatory coupling space to compute the gene-gene relationship matrix (*ϕ*). **(b)** Greedy insertion rules for gene hierarchy construction. For each gene, its placement relative to an existing node is determined by topological rules (R1–R4). **(c)** The resulting gene hierarchy enables multi-resolution cell clustering.

A reliable cell-type partition is not an observation but a construct, supported by a set of genes whose expression patterns consistently align with the proposed grouping[10, 11]. The credibility of a partition increases with both the number and strength of supporting genes, and importantly, these genes mutually support one another. Conventional methods like PCA model gene relationships via global correlation, assessing whether two genes are consistently high or low across all cells. However, Pearson/Spearman correlations often fail in single-cell data, causing a large number of real gene-gene interactions to go undetected[12]. Biologically relevant coordination in cell-type definition often takes the form of Boolean regulatory logic: genes are jointly on or jointly off within specific subsets[13].

At a mathematical level, cell–cell similarity in scRNA-seq is the problem of measuring distances between gene-expression functions in a Hilbert space, and this measurement is inherently comparison-dependent rather than global[14]. Each cell can be viewed as a function *f*_*x*_ defined over genes, and any distance between two cells is obtained by applying a functional to the difference *f*_*x*_ − *f*_*y*_. Biologically, however, such functionals are never fixed across all comparisons, where different contrasts rely on different gene programs. Lineage-level distinctions (e.g. T vs. B cells) depend on one subset of genes, whereas finer intra-lineage distinctions (e.g. naïve vs. effector T cells) depend on entirely different regulatory modules [15, 16]. In Hilbert terms, each biological comparison effectively restricts evaluation to a program-specific subspace ℋ_*u*_ ⊂ ℋ, equipped with its own inner product and associated energy functional for measuring ∥ *f*_*x*_ − *f*_*y*_∥ within that subspace. Thus, the transcriptomic landscape is not a single homogeneous metric space equipped with a universal norm, but a heterogeneous family of comparison-dependent Hilbert subspaces and functionals, each selected by the particular biological question embodied in a given cell pair.

The difficulty is that realizing this comparison-dependent Hilbert structure naively destroys the geometric and statistical foundations on which current single-cell methods rely[17, 18]. If the functional used to evaluate ∥ *f*_*x*_ *− f*_*y*_∥ is allowed to vary arbitrarily with the pair (*x, y*), the resulting distances need not satisfy basic metric properties, breaking the neighborhood graphs, diffusion operators, and low-dimensional embeddings that presume a single coherent inner-product geometry. Statistically, learning a distinct functional for every cell pair is hopelessly underdetermined, as the degrees of freedom scale as 𝒪(*n*^2^) in the number of cells, far exceeding what sparse and noisy scRNA-seq measurements can support. Methodologically, existing pipelines are engineered around a fixed feature space and a global distance matrix, and even “locally adaptive” methods like PHATE[19] and UMAP[20] merely reweight coordinates or tune kernels within a shared Hilbert geometry rather than altering the geometry itself. Any viable formulation of pair-dependent distances must therefore impose a strongly structured, biologically grounded family of Hilbert subspaces and energy functionals, enabling comparison rules to vary across cell pairs without sacrificing consistency, identifiability, or computational tractability.

To resolve this geometric dilemma, we introduce GeCCo (Gene Co-expression Constructed identity), a framework that instantiates a structured family of Hilbert subspaces for robust identity construction. Rather than attempting to learn unstable distance metrics from sparse cell embeddings, GeCCo anchors cell identities within a rigorous, pre-computed hierarchy of gene programs. Specifically, we quantify genegene relationships using the *ϕ* coefficient to capture Boolean regulatory logic-distinguishing synergistic co-activation from mutual antagonism-and organize these relationships via greedy topological inference. This results in a resolution-adaptive metric where broad lineages are separated by antagonistic modules while finer subtypes are resolved by nested co-activation. We demonstrate that this gene-centric architecture yields substantially higher hierarchical consistency than cell-centric baselines on a comprehensive human immune reference and uncovers subtle transitional states in pancreatic development that standard geometric methods obscure.

## 2 Methods

### 2.1 Problem formulation: cell-cell distance in a heterogeneous transcriptomic space

Let *f*_*x*_: *G* → ℝ denote the gene expression function of a cell *x*, where *G* is the set of assayed genes. Conventional single-cell analysis assumes that all cells lie in a single ambient vector space ℝ^|*G*|^ endowed with a fixed, global metric. Under this assumption, cellcell similarity is computed by applying the same distance function to every pair of cells. However, the transcriptomic space is inherently heterogeneous: different cell pairs are distinguished by different gene programs, and thus should be compared in different feature subspaces. As a result, no fixed metric on ℝ^|*G*|^ can faithfully capture intercellular similarities across multiple biological resolutions.

To formalize this heterogeneity, we regard each cell expression profile *f*_*x*_ as an element of a function space ℋ, and define the distance between two cells *x* and *y* through a *pair-dependent Hilbert functional*. Specifically, for each cell pair (*x, y*) we associate a parameter *θ*(*x, y*), representing the gene module most relevant for comparing *x* and *y*. This parameter determines a positive-definite operator *M*_*θ*(*x,y*)_ on ℋ, which induces a Hilbert inner product

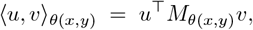

and corresponding quadratic functional

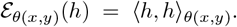

The distance between cells *x* and *y* is then defined by the energy of their expression difference:

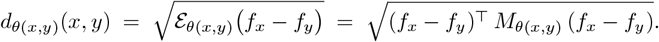

The key challenge is that allowing *M* to vary arbitrarily with (*x, y*) would destroy the global geometric consistency required for downstream analysis. We resolve this by constraining *θ*(*x, y*) to range over the nodes of a gene-program hierarchy, which provides a structured, finite, and biologically grounded family of Hilbert subspaces. Each *M*_*θ*_ is the inner-product operator associated with the corresponding gene module, and the hierarchy guarantees compatibility among these operators across scales. This framework defines a mathematically coherent and biologically meaningful family of pair-dependent metrics, enabling cell identities to be constructed from gene-program structure rather than global transcriptomic geometry.

### 2.2 Quantification of Boolean Regulatory Coupling

To capture the logic of gene regulation, which frequently operates in sharp on-off states rather than continuous gradients, we project the continuous transcriptomic manifold onto a Boolean hypercube. Let 𝒳 ∈ ℝ ^*G*×*N*^ be the normalized expression matrix. We define a binary state matrix **B** ∈ {0, 1}^*G×N*^ via an indicator function 𝕀 (·), where *B*_*gi*_ = 1 if *X*_*gi*_ *> τ* (with *τ* = 0.5 TPM) and 0 otherwise. We model the expression state of each gene *g* as a Bernoulli random variable *I*_*g*_ with success probability 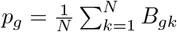.

To quantify the coupling strength between any gene pair (*i, j*), we compute the *ϕ* coefficient, which is mathematically equivalent to the Pearson correlation coefficient for two binary variables. Instead of traditional contingency tables, we formulate *ϕ*_*ij*_ directly via the centered covariance of the observed boolean vectors:

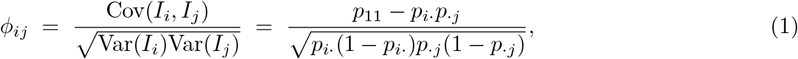

where *p*_*i•*_ and *p •* _*j*_ denote the marginal probabilities of gene activation, and *p*_11_ denotes the joint probability of co-activation (estimated as 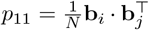, where **b**_*i*_ is the *i*-th row of **B**). This formulation captures the standardized linear dependence between regulatory states: *ϕ*_*ij*_ *>* 0 implies synergistic co-activation, while *ϕ*_*ij*_ *<* 0 indicates mutual exclusivity.

The statistical significance of these directional associations was rigorously assessed using the hypergeometric distribution (one-tailed Fisher’s exact test). Specifically, for *ϕ*_*ij*_ *>* 0, we tested the null hypothesis that gene *i* and *j* are activated independently against the alternative of positive dependence (Odds Ratio *>* 1); conversely, for *ϕ*_*ij*_ *<* 0, we tested against negative dependence (Odds Ratio *<* 1). To control for false discoveries in the high-dimensional gene pair space, *p*-values were adjusted using the Benjamini-Hochberg procedure. Valid regulatory edges were retained only if they satisfied a dual threshold of statistical significance (FDR ≤ 0.05), ensuring the reconstructed hierarchy is grounded in robust non-random associations.

### 2.3 Hierarchical gene module construction for cell pair-dependent metric

To realize a pair-dependent metric as a family of Hilbert functionals, we require a structured collection of gene modules that induce compatible subspaces rather than an arbitrary partition of genes. Concretely, for each module *u* in a rooted tree 𝒯 we will later associate a Hilbert subspace ℋ_*u*_ ⊂ ℋ and a positive-definite operator *M*_*u*_ that defines the local inner product and energy functional used for comparing cells whose identities are governed by *u*. This construction imposes strong constraints on how genes can be grouped: modules must be internally coherent and externally antagonistic in a way that reflects regulatory logic and yields nested, non-contradictory subspaces.

#### Signed gene co-expression graph

Let *G* = (*V, E*^+^, *E*^−^) be a signed gene co-expression graph constructed from the *ϕ*-coefficient matrix *Φ* = (*ϕ*_*ij*_), where *V* is the set of genes, *E*^+^ = {(*i, j*): *ϕ*_*ij*_ > 0} is the set of positively correlated gene pairs, and *E*^−^ = {(*i, j*): *ϕ*_*ij*_ < 0} is the set of negatively correlated pairs. We seek a rooted tree 𝒯 = (*U, F*) together with a mapping *M*: *U* → 2^*V*^ that assigns to each node *u* ∈ *U* a gene module *M* (*u*) ⊆ *V*. The root *r* is a purely structural node with *M* (*r*) = ∅.

To ensure that the resulting hierarchy can serve as a biologically meaningful family of Hilbert subspaces, we impose three topologicalcorrelation constraints on 𝒯:

(C1) **Within-module positivity**. For any non-root node *u* and any *i, j* ∈ *M* (*u*) with *i* ≠ *j*, we require *ϕ*_*ij*_ > 0.

(C2) **Sibling antagonism**. For any two distinct child nodes *u, v* sharing the same parent *p*, and any *i* ∈ *M* (*u*), *j* ∈ *M* (*v*), we require *ϕ*_*ij*_ < 0.

(C3) **Parentchild coherence**. For any parentchild pair (*p, c*) and any *i* ∈ *M* (*p*), *j* ∈ *M* (*c*), we require *ϕ*_*ij*_ > 0.

(C1) enforces that each module encodes a positively co-regulated gene program; (C2) ensures that sibling modules represent mutually antagonistic programs that can separate lineages; and (C3) guarantees that parent modules capture broad programs that remain positively coupled to all their downstream refinements. These constraints are precisely what is needed to interpret each *M* (*u*) as a coherent Hilbert subspace and to maintain compatibility among the subspaces across the hierarchy.

#### Greedy topological construction

The construction of 𝒯 is guided by the biological intuition that hierarchy emerges from the resolution of conflict. Broad lineages (higher nodes) are defined by genes that are broadly co-expressed (synergistic) across many subtypes, whereas specific subtypes (leaves) are distinguished by genes that are mutually antagonistic.

To operationalize this intuition, we construct the tree 𝒯 via a greedy hierarchical algorithm that incrementally inserts genes into modules while preserving (C1)(C3). The root node *r* is initialized with *M* (*r*) = ∅.

##### Initialization via an anchor gene

Instead of selecting an arbitrary correlated pair, we first identify an anchor gene *a* as the gene with the largest connectivity after statistical testing, i.e., the gene with the maximal number (or total weight) of significant regulatory edges in *E*. Biologically, this anchor represents a highly connected and globally influential gene in the co-expression graph.

We then identify (i) the gene *b*^+^ that is most positively correlated with *a*, defined as *b*^+^ = arg max_*j* ∈ *V*\_ {_*a*_} *ϕ*_*aj*_, and (ii) the gene *b*^−^ that is most negatively correlated with *a*, defined as *b*^−^ = arg min_*j*_ ∈_*V*_ \ {_*a*}_ *ϕ*_*aj*_.

Two initial child nodes are created under the root, with *M* (*u*^+^) = {*a, b*^+^} and *M* (*u*^−^) = {*b*^−^}, where *u*^+^, *u*^−^ ∈ *𝒞* (*r*). This initialization explicitly separates synergistic and antagonistic directions at the top level of the hierarchy, establishing an early bifurcation structure consistent with (C1)(C3).

##### Insertion order of remaining genes

The remaining genes in *V* \ {*a, b*^+^, *b*^−^} are inserted one by one. The insertion order is determined by decreasing prevalence across cells: genes are sorted by the number of cells in which they are expressed (nonzero or above threshold), from highest to lowest.

This design reflects the biological assumption that genes expressed in many cells are more likely to represent housekeeping or broadly shared programs, whereas genes expressed in fewer cells are more likely to act as subtype-specific markers. Thus, broadly expressed genes are inserted earlier in the hierarchy, and more specific genes are inserted later, allowing the tree to gradually refine into subtype-level branches.

##### Adaptive correlation threshold

To mitigate potential confounding between subtype-specific genes and broadly expressed (housekeeping) genes-where subtype genes may appear spuriously correlated with dominant global programs-we introduce an adaptive correlation threshold that depends on the insertion stage.

Suppose there are *n* genes in total, and let *g*_*k*_ denote the *k*-th gene inserted (according to the above order). We define the stage-dependent threshold as 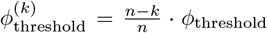, where *ϕ*_threshold_ is a base threshold parameter.

Thus, earlier-inserted genes (small *k*) are subject to stricter correlation constraints, whereas later-inserted genes (large *k*) are evaluated under a more permissive threshold. This adaptive mechanism ensures that early high-prevalence genes establish robust, globally consistent modules, while later low-prevalence genes can flexibly specialize within finer-grained substructures.

##### Incremental top-down insertion

For each gene *g* to be inserted, we perform a top-down traversal starting from the root. At a current internal node *u*, with children 𝒞 (*u*), we classify each child module *c* ∈ 𝒞 (*u*) according to the sign pattern of *ϕ*_*gj*_ for *j* ∈ *M* (*c*):

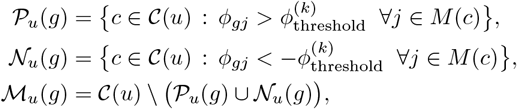

corresponding respectively to purely positive, purely negative, and mixed-sign relationships between *g* and the genes in each child module under the stage-adaptive threshold.

Placement decisions at node *u* follow four biologically motivated rules that preserve (C1)(C3).

**R1 (absorption into a unique positive child)**. If 𝒫_*u*_(*g*) contains exactly one module *c* and ℳ_*u*_(*g*) is empty, then *g* is recursively inserted into the subtree rooted at *c*. When *c* is a leaf, we update *M* (*c*) ← *M* (*c*) ∪ {*g*}, preserving (C1) by construction.

- **R2 (creation of an intermediate parent)**. If 𝒫 _*u*_(*g*) contains at least two modules and ℳ_*u*_(*g*) is empty, then *g* is positively correlated with multiple siblings. In this case we create a new internal node *v* as a child of *u* with module *M* (*v*) = {*g*}, and reattach all modules in 𝒫_*u*_(*g*) as children of *v*. This preserves (C2) among the reattached siblings and enforces (C3) between *v* and its descendants.

**R3 (Creation of a new sibling lineage)**. If 𝒫_*u*_(*g*) is empty and 𝒩_*u*_(*g*) comprises all children of *u*, gene *g* exhibits mutual exclusivity with every existing module at the current level. This pattern suggests that *g* characterizes a novel biological program distinct from the existing lineages. Therefore, we create a new child node *v* with *M* (*v*) = {*g*} and attach it as a sibling under *u*. This placement satisfies (C2) by construction, as *g* is antagonistic to all its new siblings.

- **R4 (conflicts at leaves)**. If a leaf child *c* belongs to ℳ_*u*_(*g*) (mixed signs within *M* (*c*)), we resolve the conflict by bifurcating *c* into two new child modules *c*^+^ and *c*^−^ under *u*, whose gene sets collect the positively and negatively correlated subsets with respect to *g*. The original module *c* is removed, and *g* is then inserted recursively under the appropriate new child or excluded if constraints cannot be satisfied.

The traversal for gene *g* terminates when *g* is either absorbed into a leaf module (R1), becomes a new internal module (R2), or is deemed incompatible and excluded (R3/R4 after exhaustive backtracking). The algorithm iterates over all genes in *V*, resulting in a signed hierarchical tree 𝒯 that satisfies (C1)(C3) by construction. In the worst case, the complexity is 𝒪 (|*V*| ^2^), dominated by the evaluation of *ϕ*_*gj*_ across genes during insertion.

#### From modules to Hilbert subspaces

Given the resulting tree 𝒯, each node *u* defines a gene index set *M* (*u*) and hence a finite-dimensional Hilbert subspace

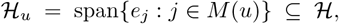

where {*e*_*j*_} is the canonical basis of gene-wise expression functions. Because of (C1)(C3), these subspaces are nested along root-to-leaf paths and separated by antagonistic relationships across siblings, which allows us to attach a positive-definite operator *M*_*u*_ to each node and interpret

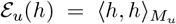

as a coherent energy functional for measuring expression differences restricted to the program encoded by *u*. In later sections, we use these node-specific functionals to define pair-dependent distances: for a given pair of cells (*x, y*), the parameter *θ*(*x, y*) selects an appropriate node *u* in 𝒯, and the distance *d*_*θ*(*x,y*)_(*x, y*) is computed as the norm of *f*_*x*_ − *f*_*y*_ in the Hilbert subspace ℋ_*u*_.

### 2.4 Cell-to-module assignment via hierarchical gene activation

Given a signed gene module tree 𝒯 as described in Section 2.3, we next assign each cell to a unique node in 𝒯 so that its identity is anchored in a specific gene-program subspace. This step serves two purposes: biologically, it maps cells onto a hierarchy of gene programs that encode increasingly specific regulatory states; geometrically, it provides a discrete index on 𝒯 that will later be used to select the Hilbert subspace and energy functional for defining pair-dependent distances between cells.

#### Module activity scores

Let *X* ∈ ℝ^*n×p*^ denote the normalized expression matrix, where *X*_*ig*_ is the expression of gene *g* in cell *i*. For each gene *g*, we compute a standardized expression

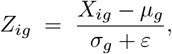

where *µ*_*g*_ and *σ*_*g*_ are the mean and standard deviation of *X·*_*g*_ across all cells, and *ε* is a small constant to stabilize low-variance genes. For each node *u* ∈ 𝒯 with gene set *G*_*u*_, we define the activation score of module *u* in cell *i* as a robust aggregate of standardized expression:

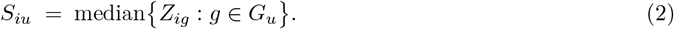

This score approximates the projection of *f*_*i*_ onto the subspace ℋ_*u*_ spanned by *G*_*u*_, while being robust to outliers within the module. Because modules on any roottoleaf path are nested and siblings are antagonistic, the collection {*S*_*iu*_: *u* ∈ 𝒯} encodes a structured profile of gene-program activation for cell *i* across biological scales.

#### Top-down hierarchical assignment

To assign cell *i* to a node in 𝒯, we perform a deterministic top-down traversal that follows the strongest activated branch while enforcing both relative and absolute specificity. Let *u*_0_ denote the root. Starting from *u*_0_, at a generic node *u* with children 𝒞 (*u*) we identify the child with maximal activation

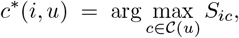

and define the corresponding maximum score

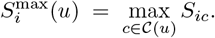

We proceed from *u* to *c*^*∗*^(*i, u*) only if two criteria are satisfied:

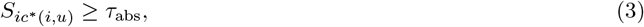

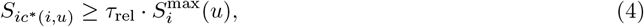

where *τ*_abs_ controls an absolute activation threshold and *τ*_rel_ ∈ (0, 1] enforces that the chosen child is sufficiently dominant relative to its siblings. If either condition fails at node *u*, the traversal for cell *i* terminates and *i* is assigned to *u* as its most specific reliable module. If both conditions hold and *c*^*∗*^(*i, u*) is non-leaf, we update *u* ← *c*^*∗*^(*i, u*) and repeat the procedure; if *c*^*∗*^(*i, u*) is a leaf, we assign *i* to this leaf.

Formally, the assignment function *a*: {1, …, *n*} → *U* is defined by

*a*(*i*) = the terminal node reached by iterating *u*_*t*+1_ = *c*^*∗*^(*i, u*_*t*_) under conditions (3)–(4), starting at *u*_0_.

Because each step follows a unique maximizer and the tree has no cycles, every cell is assigned to exactly one node in 𝒯.

##### Algorithm 1: Construct signed co-expression module tree from signed gene network *G* = (*V, E*^+^, *E*^−^).

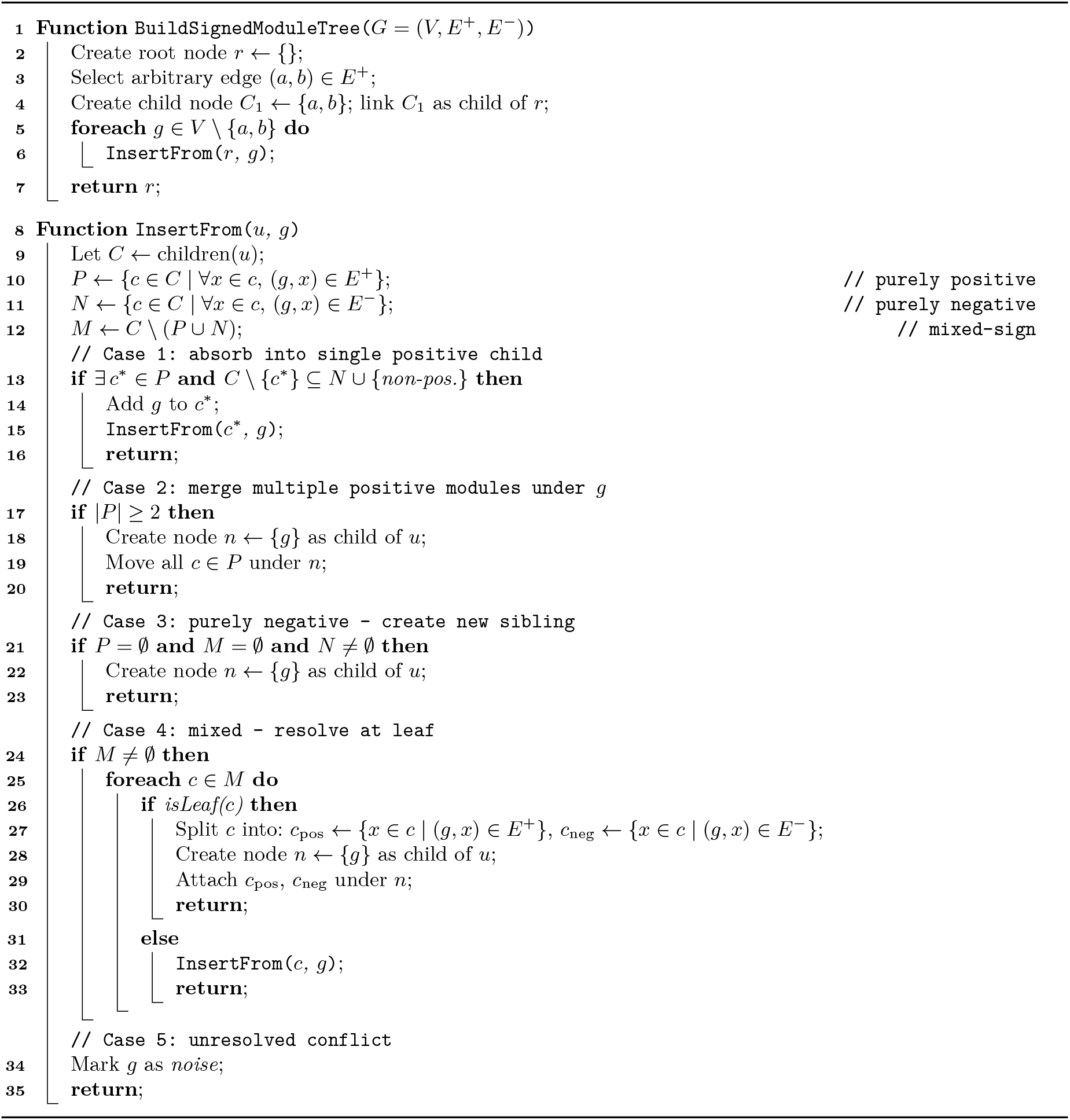

#### Interpretation and role in the metric

This hierarchical assignment yields three biologically interpretable outcomes. Cells assigned to leaf nodes correspond to pure transcriptional states dominated by a single, highly specific gene program. Cells assigned to internal nodes represent transitional or hybrid states, for which no single descendant program is sufficiently dominant relative to its siblings. Cells that fail to satisfy the absolute threshold at any non-root node can be flagged as weakly activated or potentially artifactual and optionally excluded from downstream analyses.

Crucially, the assignment *a*(*i*) also supplies the discrete structural index needed for our pair-dependent metric. For a pair of cells (*i, j*), the parameter *θ*(*i, j*) used in the Hilbert functional is derived from their positions on 𝒯, for example via the lowest common ancestor of *a*(*i*) and *a*(*j*) or another biologically motivated rule. This ensures that expression differences between *i* and *j* are measured in the gene-program subspace that is most appropriate for that pair, rather than in a fixed global feature space.

## 3 Results

### 3.1 GeCCo demonstrates superior hierarchical consistency and robustness

To evaluate hierarchical consistency in the absence of absolute ground truth, we benchmarked GeCCo on the human Bone Marrow Mononuclear Cell (BMMC) reference atlas[21]. We compared performance against a diverse array of representative clustering algorithms, including Scanpy[8], Cytocipher[22], SC3[23]and sc-SHC[24].

As shown in **Figure 3a**, GeCCo demonstrated superior performance, achieving the highest median Adjusted Rand Index (ARI) in local consistency evaluations while maintaining a robust global consistency score. The baseline methods exhibited distinct failure modes rooted in their design assumptions. Graph-based and consensus methods (e.g., Scanpy, SC3) enforce a uniform resolution over a density-varying manifold; constrained by a static global feature space, they inevitably suppress local granularity to preserve global topology. Conversely, recursive approaches (sc-SHC) treat hierarchical splits as independent statistical events. Lacking global regularization, this greedy strategy creates irreversible fragmentation, failing to reconstruct the coherent lineage structure captured by GeCCo.

**Fig. 3.**
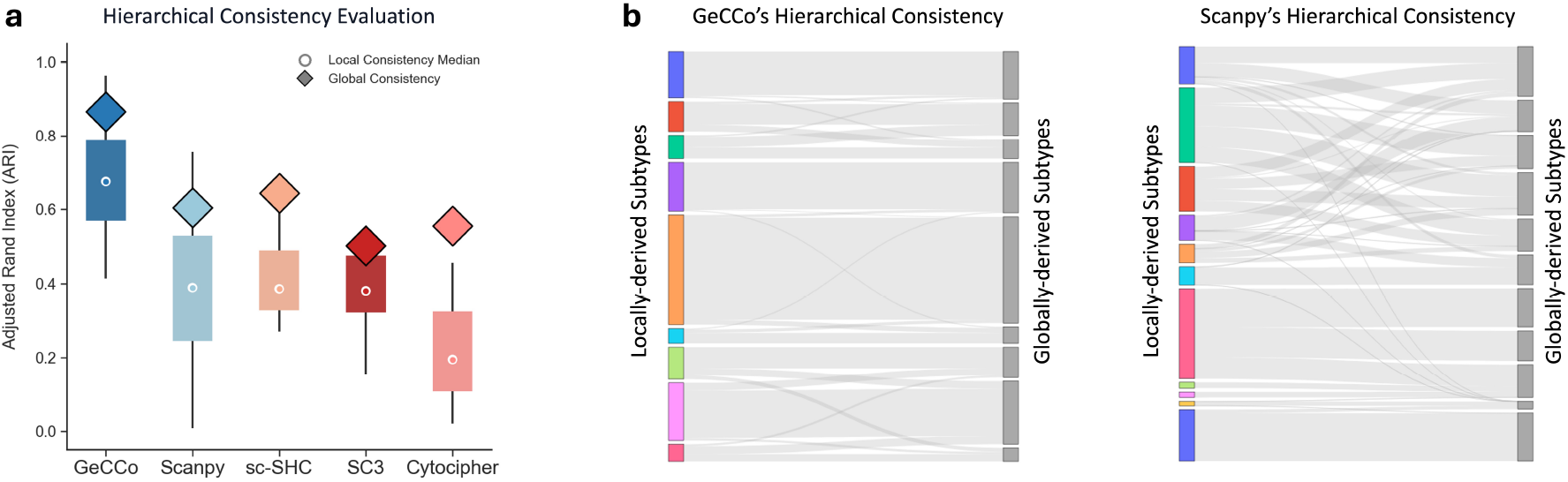
Evaluation of hierarchical consistency and clustering stability. **(a)** Quantitative comparison of GeCCo against baseline methods using ARI. Diamonds represent the global consistency score (alignment between the global clustering result and the aggregated local sub-clustering results). Boxplots display the distribution of local consistency scores across individual subtypes, with white dots indicating the median. **(b)** Sankey diagrams visualizing the structural correspondence between locally derived subtypes (left column) and globally derived subtypes (right column). GeCCo exhibits minimal errors in major branch splits and only misclassifies regions with subtle differences, whereas entangled flows in Scanpy indicate some major branch splits.

To dissect the structural basis of these inconsistencies, we visualized the mapping between locally derived and globally derived subtypes (Figure 3b). GeCCo revealed clean, parallel transitions, indicating that its global lineage definitions are structurally compatible with local subtype distinctions. In contrast, competitor methods exhibited chaotic, crisscrossing flows-a visual signature of hierarchical discordance-where cells identified as a homogeneous group in one context were fragmented into conflicting clusters in another. This structural instability is further corroborated by the embedding manifolds in Supplementary Figure 3, where GeCCo alone preserves the topological alignment between global clusters and local substructures.

We attribute the hierarchical inconsistency in baseline methods to their reliance on global feature selection. Standard pipelines prioritize Highly Variable Genes (HVGs) calculated across the entire dataset, a strategy that systematically masks genes critical for local distinctions but lacking global variance. By shifting from variance-based selection to co-expression-based construction, GeCCo resolves the tension between global topology and local heterogeneity, ensuring that identified substates represent biologically meaningful programs rather than computational artifacts of the analysis scope.

### 3.2 GeCCo resolves heterogeneity in endocrine progenitors and identifies a concentrated division phase

We applied GeCCo to the real mouse pancreas dataset[25], specifically zooming in on the Ngn3-high endocrine progenitor (EP) population (Figure 4a). While often considered a homogeneous transient cluster in conventional analyses, GeCCo uncovered a structured underlying heterogeneity driven by three gene modules (GM1, GM2, and GM3). The Phi-score heatmap (Figure 4b) reveals a striking pattern of transcriptional antagonism: genes within each module are co-expressed (red blocks), while exhibiting strong mutual exclusivity against genes in other modules (blue off-diagonal blocks).

**Fig. 4.**
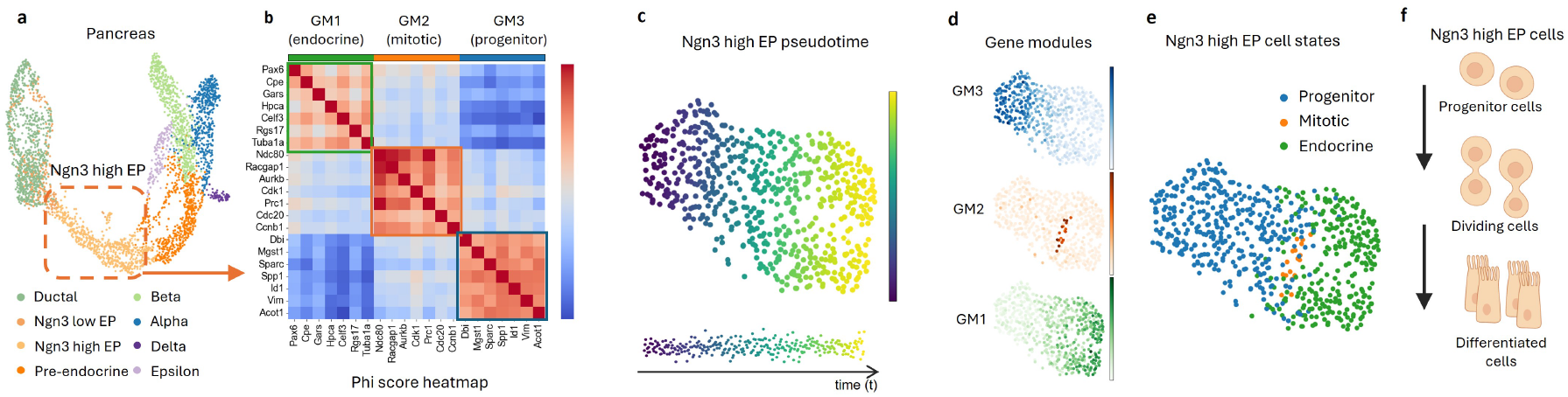
GeCCo unravels three substates in Ngn3-high endocrine progenitors, implying a division phase prior to fate commitment.

Biological annotation of these modules defines three distinct functional states: GM3 (Progenitor) marks the early uncommitted state; GM1 (Endocrine) represents the differentiated outcome; and notably, GM2 (Mitotic) captures a highly proliferative state enriched with cell cycle regulators.

To reconstruct the developmental progression, we aligned these cells along a pseudotime trajectory (Figure 4c), which establishes a general temporal direction from early progenitors to differentiated cells. Projecting the GeCCo modules onto this trajectory reveals a clear sequential transition (Figure 4d). When cells are classified by their dominant module (Figure 4e), a distinct “sandwich” structure emerges: the Mitotic state (GM2, orange) is spatially and temporally located strictly between the Progenitor state (GM3, blue) and the Endocrine state (GM1, green).

This observation leads to a crucial biological insight summarized in our model (Figure 4f). The exclusive presence of GM2 as a bridge between GM3 and GM1 suggests that differentiation in Ngn3-high cells is not a continuous, asynchronous drift. Instead, cells likely undergo a concentrated division phase: a synchronized entry into the cell cycle to expand the progenitor pool before collectively exiting to commit to the endocrine fate. GeCCos ability to leverage gene antagonism was essential to resolving this transient but mandatory intermediate state, which is easily obscured in standard clustering analyses.

## 4 Discussion

Defining cell identity requires measuring similarity over context-dependent gene subsets rather than a fixed feature space. GeCCo addresses this by constructing identities within a structured family of Hilbert sub-spaces, ensuring that the distance metric adapts to biological resolution without breaking global geometric consistency. By resolving conflicts between synergistic and antagonistic gene programs, GeCCo aligns local adaptivity with global hierarchical consistency-a critical property often lacking in standard cell-centric pipelines.

A key insight of our framework is that identity is defined not just by expression, but by systematic exclusion. Unlike methods relying solely on positive correlation, GeCCo leverages gene antagonism to strictly delineate transitional states. This capability was crucial in uncovering the hidden “mitotic bridge” in pancreatic progenitors, demonstrating that biological granularity often resides in the negative space of gene regulation that standard variance-based methods discard.

GeCCo operates under specific assumptions. First, modeling regulation via pairwise *ϕ* coefficients simplifies continuous kinetics and may miss high-order combinatorial logic (e.g., multi-gene XOR). Second, the strict tree topology, while ideal for lineage differentiation, may fail to effectively model cyclic or convergent trajectories. Finally, constructing global co-expression networks entails higher computational costs (𝒪|*G*^2^|) than standard feature selection, necessitating optimization for massive datasets.

As single-cell atlases grow, GeCCo offers a stable foundation where identities are defined by enacted gene programs rather than dataset-specific embeddings. This shift from ad hoc clustering to programmatic cell typing paves the way for universally consistent reference atlases.GeCCo is publicly available at https://github.com/Lu7-ydd/gecco.

## Acknowledgements

This work is supported by STI2030-Major Projects 2022ZD0212400, Lingang Laboratory grant LGL-8888, STCSM grant 24510714300 and 20DZ2254400, GuangDong Basic and Applied Basic Research Foundation grant 2023B1515120006 and SJTU Science-Medicine interdisciplinary grant YG2026ZD09 and 24X010301456.

