## Supplementary for "Gene-First Identity Construction for Robust Cell Identification in Single-Cell Transcriptomics" (from another version of this preprint: version 1, 2026-02-26)

### GeCCo supplement

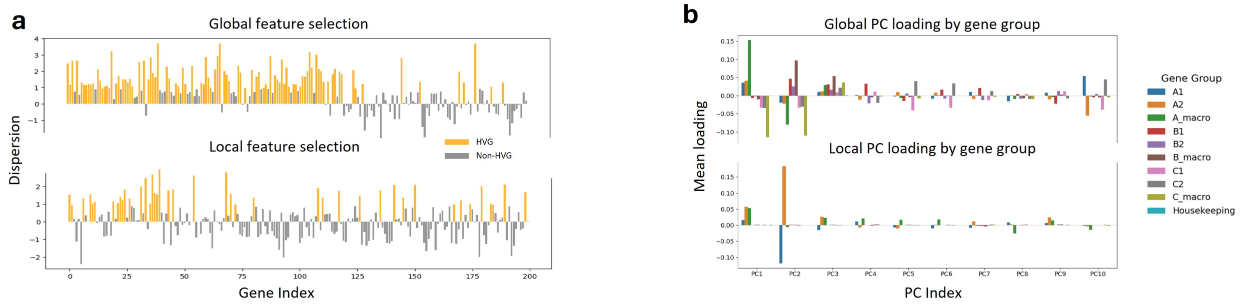


**Fig. S1. Mechanistic basis of global context bias in feature selection and dimensionality reduction.** This figure provides the underlying explanation for the hierarchical inconsistency observed in Figure 1. **(a)** Comparison of feature selection (dispersion) in global versus local contexts. Genes that drive local subtype distinctions (identified as highly variable genes, HVG, in orange in the local context, bottom panel) are frequently masked due to lower relative dispersion when calculated globally across the entire dataset (grey bars in the top panel). **(b)** Contrast in mean Principal Component (PC) loadings by gene group between global and local analyses. Global PCs (top panel) are dominated by loadings from macro-lineage markers (e.g., A__macro, B__macro), failing to capture variance from subtype-specific drivers. In contrast, local PCA (bottom panel, exemplified using lineage A) identifies distinct axes of variation heavily loaded by subtype markers (e.g., A1, A2), which are necessary to resolve fine-grained substates.

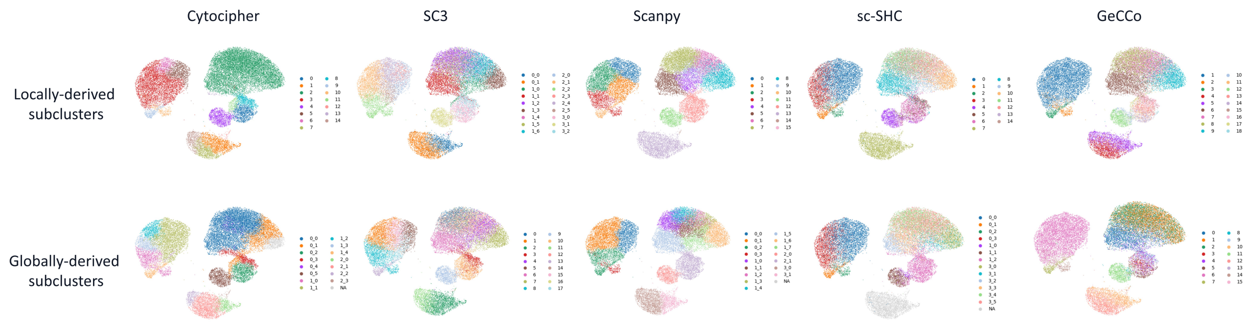


**Fig. S3. Qualitative comparison of hierarchical consistency across clustering algorithms on the BMMC dataset.** The figure visually assesses the stability of cell identity definitions when analysis scope shifts from local to global. **Top row (Locally-derived subclusters):** UMAP embeddings colored by subtype labels obtained by independently sub-clustering isolated lineages. This serves as the high-resolution biological reference. **Bottom row (Globally-derived subclusters):** UMAP embeddings of the same cells colored by cluster labels derived from clustering the entire dataset directly. **Comparison:** A robust method should yield global clusters (bottom) that spatially correspond to local subtypes (top). Baseline methods (Cytocipher, SC3, Scanpy, sc-SHC) exhibit visible structural discordance, where biologically distinct local subtypes are frequently merged or fragmented in the global view. In contrast, **GeCCo** demonstrates superior topological consistency, maintaining clear separation of subtypes even within the global embedding, thereby minimizing the discrepancy between local and global identity definitions.
